# Long-read Transcriptomics of Caviid Gammaherpesvirus 1: Compiling a Comprehensive RNA Atlas

**DOI:** 10.1101/2024.12.11.627975

**Authors:** Gábor Torma, Ákos Dörmő, Ádám Fülöp, Dóra Tombácz, Máté Mizik, Amanda M. Pretory, See-Chi Lee, Zsolt Toth, Zsolt Boldogkői

## Abstract

Caviid gammaherpesvirus 1 (CaGHV-1), formerly known as the guinea pig herpes-like virus, is an oncogenic gammaherpesvirus with a sequenced genome but an as-yet uncharacterized transcriptome. Using nanopore long-read RNA sequencing, we annotated the CaGHV-1 genome and constructed a detailed transcriptomic atlas. Our findings reveal diverse viral mRNAs and non-coding RNAs, along with mapped promoter elements for each viral gene. We demonstrated that the CaGHV-1 RTA lytic cycle transcription factor activates its own promoter, similar to KSHV, and that the CaGHV-1 ORF50 promoter responds to RTA proteins from other gammaherpesviruses, highlighting the evolutionary conservation of RTA-mediated transcriptional mechanisms. Additionally, our analysis uncovered extensive transcriptional overlap within the viral genome, suggesting a role in regulating global gene expression. Given its tumorigenic properties, broad host range, and non-human pathogenicity, this work establishes CaGHV-1 as a promising small animal model for investigating human gammaherpesvirus pathogenesis.

**IMPORTANCE:** The molecular underpinnings of gammaherpesvirus pathogenesis remain poorly understood, partly due to limited animal models. This study provides the first comprehensive transcriptomic atlas of CaGHV-1, highlighting both coding and non-coding RNAs and revealing regulatory elements that drive viral gene expression. Functional studies of the CaGHV-1 RTA transcription factor demonstrated its ability to self-activate and cross-activate promoters from homologous gammaherpesviruses, reflecting conserved mechanisms of transcriptional control. These findings solidify CaGHV-1 as a unique and versatile small animal model, offering new opportunities to investigate gammaherpesvirus replication, transcriptional regulation, and tumorigenesis in a controlled experimental system.

## INTRODUCTION

Gammaherpesviruses are a subfamily of herpesviruses that establish lifelong latency in their hosts by infecting lymphocytes. They are medically significant due to their association with various cancers, including lymphomas as well as endothelial and epithelial tumors (1, 2). Epstein-Barr virus (EBV), one of the most well-known human gammaherpesviruses, is associated with infectious mononucleosis and various malignancies (3). Another human gammaherpesvirus is Kaposi’s sarcoma-associated herpesvirus (KSHV), responsible for Kaposi’s sarcoma, primary effusion lymphoma, and a subset of multicentric Castleman diseases, particularly in immunocompromised individuals such as those with AIDS (4). Current research predominantly relies on the murine model, specifically the murine gammaherpesvirus 4 strain 68 (MHV68), for studying the pathogenesis of human gammaherpesviruses *in vivo* (5). While MHV68 has been valuable for understanding the key viral and host determinants of gammaherpesvirus infection, it is constrained by its limited sequence homology and significant physiological differences compared to human gammaherpesviruses (6, 7). Non-human primate models, such as those employing Rhesus rhadinovirus (RRV) and Retroperitoneal fibromatosis-associated herpesvirus (RFHV), offer more relevant biological insights into human gammaherpesvirus pathogenesis, but their use is limited due to the expenses of maintaining the animals, ethical concerns, the need for specialized facilities, and the complexity of conducting long-term studies in these models (8, 9). These limitations highlight the need for an alternative, affordable small animal model to study human gammaherpesvirus infection and pathogenesis.

Caviid gammaherpesvirus 1 (CaGHV-1), first identified in 1969, was recently sequenced and subsequently classified as a rhadinovirus within the gammaherpesvirus subfamily (10, 11). Previously, CaGHV-1 was referred to as the guinea pig herpes-like virus (GPHLV); however, this term is misleading, as it refers to an actual herpesvirus rather than a "herpes-like" virus. To address this, Stanfield and colleagues proposed renaming it Caviid gammaherpesvirus 1 (11). This revised name will be used consistently throughout the manuscript. The genome of CaGHV-1 spans 103,374 base pairs, with a GC content of 35.4% (11). It encodes 75 predicted open reading frames (ORFs), the majority of which are homologous to genes found in human gammaherpesviruses, such as EBV and KSHV. However, detailed transcriptomic data and the precise mapping of the viral genes are still lacking.

The introduction of long-read RNA sequencing (lrRNA-Seq) technology has revolutionized viral transcriptomics, providing a more comprehensive and accurate mapping of viral RNAs. This approach has led to the discovery of previously unidentified RNA molecules and offered deeper insights into gene expression patterns and regulatory mechanisms (12). It is increasingly evident that viral transcriptomes exhibit a complexity far beyond what was previously understood (13–16). The transcriptomes of several herpesviruses (17–20), including gammaherpesviruses (21–23), have been uncovered using this technique alone or in combination with short-read sequencing.

In this study, we present a comprehensive transcriptomic analysis of CaGHV-1, including the identification of cis-regulatory elements that may control the expression of viral genes. Additionally, we report the first functional evaluation of the transcriptional activity of the CaGHV-1 ORF50-encoded protein, which shares homology with RTA, the lytic cycle inducer in gammaherpesviruses.

## RESULTS

### General attributes of the transcriptomic analysis of CaGHV-1

To characterize the CaGHV-1 transcriptome, we analyzed the poly(A)-selected RNA fraction from infected cells using direct RNA sequencing (dRNA-Seq) and direct cDNA sequencing (dcDNA-Seq) on the ONT PromethION platform. The sequencing reads were aligned to the viral genome (OQ679822.1) using the minimap2 software. For transcriptome annotation, we utilized the LoRTIA toolkit developed in our laboratory (24). The dcDNA-Seq samples yielded a total of 71,651,155 reads, of which 5,260,508 were identified as viral reads. The dRNA-Seq generated 9,833,670 reads, which included 1,208,372 viral reads. The average read length of the dcDNA-Seq reads was 575.33 nt, while the dRNA-Seq reads had an average length of 1,013.17 nt (**Supplemental Table 1; Figures 1 A-C**). The LoRTIA software validates the quality of poly(A) sequences and sequencing adapters, while filtering out incorrect transcription start sites (TSSs), transcription end sites (TESs), and introns arising from RNA degradation, erroneous reverse transcription, artefactual PCR amplification, or sequencing mispriming. To enhance the accuracy of transcript annotations generated by the LoRTIA program, stricter filtering criteria were implemented: TSSs and TESs were considered valid only if supported by at least three dcDNA-Seq samples and one dRNA-Seq sample, while introns were identified solely based on dRNA-Seq data and subsequently verified using dcDNA-Seq data. In this study, we present only the canonical transcripts, totaling 278 (**Figure 1D**), with an average length of 4283.104 nt.

**Figure 1.**
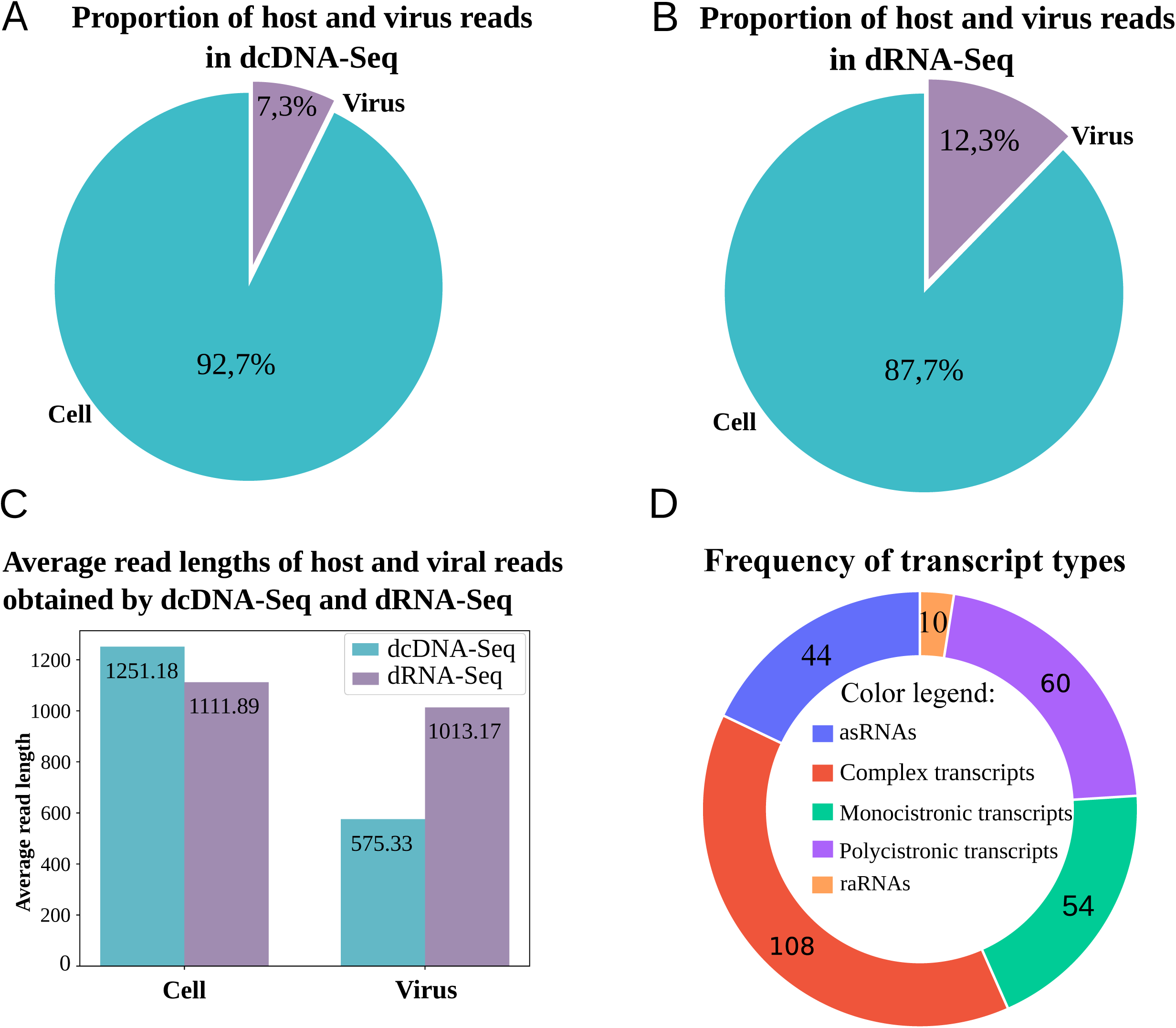
Coverage and average read lengths generated by the ONT-PromethION LRS platform. **(A)** dcDNA-Seq sequencing illustrates the proportion of host/virus reads, while **(B)** presents the proportion of host /virus reads for dRNA-Seq sequencing. **(C)** The histogram depicts the distribution of read lengths obtained from dcDNA-Seq and dRNA-Seq in host cell and virus samples. **(D)** Transcript types.

### Identification of promoters and transcription start sites of the virus

The initiator sequence at TSSs (Py A N U/A) in the CaGHV-1 genome was found to exhibit lower conservation compared to standard eukaryotic sequences (25). Notably, G/A nucleotides predominated at the TSS, with G nucleotides being especially frequent at the +1 position and T/C nucleotides primarily occupying the −1 position preceding the TSS (**Figure 2A**). This enrichment of G nucleotides has also been observed in the initiator element (Inr) of the VP5 promoter in herpes simplex virus type 1 (HSV-1) (26–28). Additionally, this Inr motif was identified in several other herpesviruses, including Epstein-Barr virus (EBV) and bovine herpesvirus 1 (BoHV-1). This enrichment of G nucleotides has also been observed in the initiator element (Inr) of the VP5 promoter in herpes simplex virus type 1 (HSV-1) (27) and identified in other herpesviruses, including Epstein-Barr virus (EBV), bovine herpesvirus 1 (BoHV-1) (22, 26, 28). We identified 92 potential TATA boxes in the CaGHV-1 genome, with an average distance of 31.80 nucleotides upstream of the TSSs. Additionally, we found 18 putative CAAT boxes, averaging 112.33 nucleotides upstream, and 5 potential GC boxes approximately 49.2 nucleotides upstream of the TSSs (**Figures 2B** and **2C**). Promoter elements within the −20 to -40 region are notably enriched in T/A nucleotides (**Supplemental Table 2**). Most of these promoter elements contain a TATA box sequence with the TATTWAA motif, which was previously detected in KSHV (29). This motif plays a key role in initiating the transcription of late genes. This analysis led to the annotation of 162 canonical transcripts. The TSS corresponded to the PAN non-coding RNA (ncRNA), supported by 1,661,506 reads, indicating an exceptionally high transcriptional state. The second most abundant TSS was associated with ORF67, with 65,034 reads. Several viral transcripts, such as the mRNAs of ORF75, ORF52, ORF59, ORF25, and ORF26, also displayed a substantial number of transcript ends.

**Figure 2.**
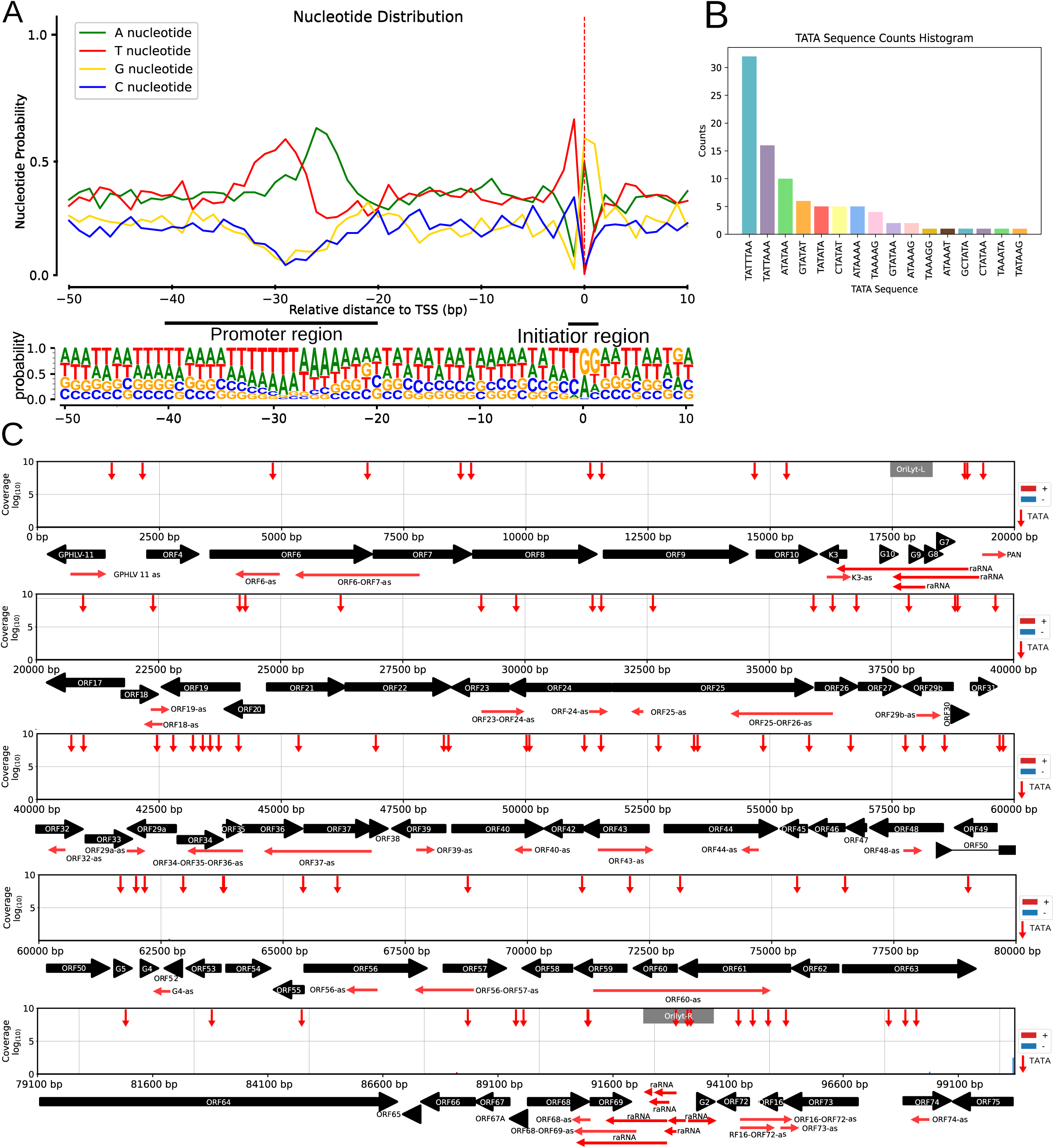
Characterization and distribution of the 5’ -ends of CaGHV-1 transcripts. **(A)** The graph illustrates the probability distribution of nucleotides within the −50 to +10 base pair interval surrounding the transcription start sites (TSSs). For most TSSs, G/A nucleotides are preferred at the +1 and +2 positions, while C/T nucleotides are favored at the −1 position. **(B)** Showing the distribution of canonical eukaryotic TATA boxes in the TSSs detected by LoRTIA. **(C)** The gray lines depict the distribution of TSSs along the viral genome as detected by LoRTIA, with the line sizes representing the TSS counts on a logarithmic scale (Log₁₀). Red vertical arrows represent the annotated TATA boxes, black horizontal arrows indicate the ORFs of genes, and green horizontal arrows represent replication origin-associated RNA and antisense RNA molecules.

### Identification of polyadenylation signals and transcription end sites of the virus

In this work, we annotated 140 canonical TESs, 131 of which were associated with polyadenylation signals (PAS), with an average distance of 25.94 nucleotides between the TESs and their corresponding poly(A) signals (**Supplemental Table 3**). The TES environment is defined by A/C cleavage sites and a preference for U/G-rich downstream elements, consistent with eukaryotic transcription termination consensus sequences. Notably, poly(A) signals are predominantly located within the 50-nucleotide upstream region (**Figures 3A** and **B**). Furthermore, we mapped TES positions across the entire viral genome using dcDNA-seq, which was validated with dRNA-Seq (**Figure 3C)**.

**Figure 3.**
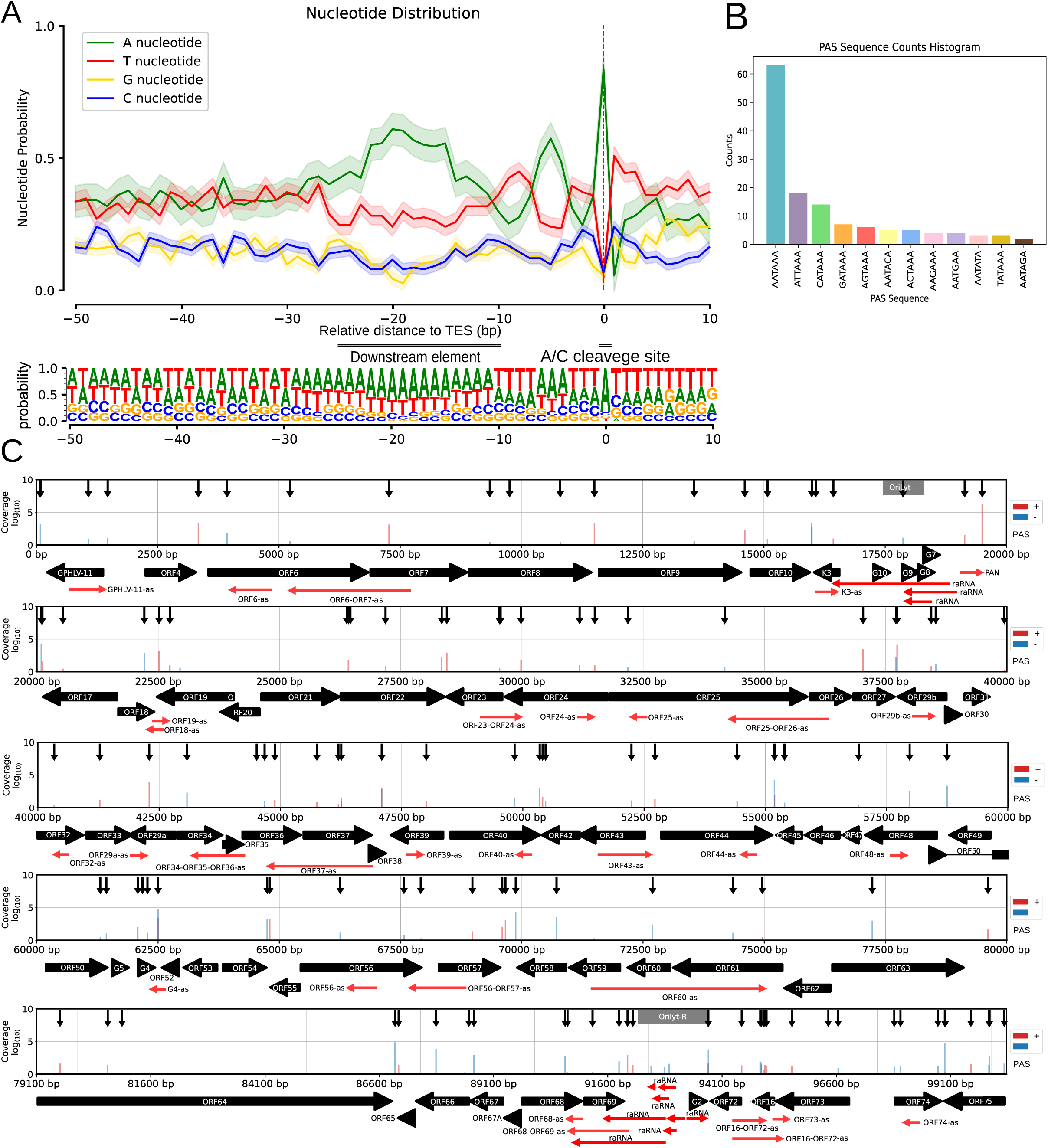
Characterization and distribution of the 3’-ends of CaGHV-1 transcripts. **(A)** The nucleotide probability distribution within the −50 to +10 base pair region surrounding the transcription end sites (TESs). TESs are characterized by the presence of the eukaryotic A/C cleavage site and G/U-rich sequence motifs downstream. **(B)** The distribution of canonical eukaryotic TATA boxes identified in TSSs by the LoRTIA program. **(C)** The gray lines show the distribution of TSSs across the viral genome as identified by LoRTIA, with line sizes corresponding to TSS counts on a logarithmic scale (Log₁₀). Black vertical arrows mark the annotated TATA boxes, black horizontal arrows indicate gene ORFs, and green horizontal arrows represent replication origin-associated RNAs (raRNAs) as well as antisense (as) RNA molecules.

### Introns and splice junctions in the viral transcripts

To explore the splicing landscape of the viral transcriptome, we analyzed our dRNA-Seq data and identified 56 introns, all validated by dcDNA-Seq **(Supplemental Table 4)**. We annotated a higher number of spliced transcripts (79) attributed to the occurrence of alternative splicing events **(Supplemental Table 5A)**. These spliced transcripts map to several genomic regions, including GPHLV-11; ORF20-ORF21; ORF29a-ORF29b; ORF31-ORF29b; ORF38-ORF40; ORF44-ORF50 (encompassing ORFs 45-47, 48, and G3-G4); ORF54-ORF57; ORF63-ORF67; ORF72-ORF73; and ORF75 **(Figure 4)**. We note that we detected a substantially higher number (280 versus 56) of introns in the dcDNA-Seq samples, most of which are likely artifacts of cDNA sequencing methodologies, potentially caused by errors during reverse transcription or second-strand DNA synthesis. To test the reliability of our dRNA-Seq results, we conducted parallel sequencing of mpox virus (a virus lacking splicing). Our results validated the accuracy of the applied sequencing and bioinformatics methods, as no false splice sites were detected in the mpox transcripts. This conclusion was further supported by obtaining the same results with respect to the introns using both the NAGATA (30) and the LoRTIA software (24) for splicing detection. This finding suggests that all spliced transcripts, including those of low abundance, are indeed of biological origin; however, many may represent mere transcriptional noise without functional significance.

**Figure 4.**
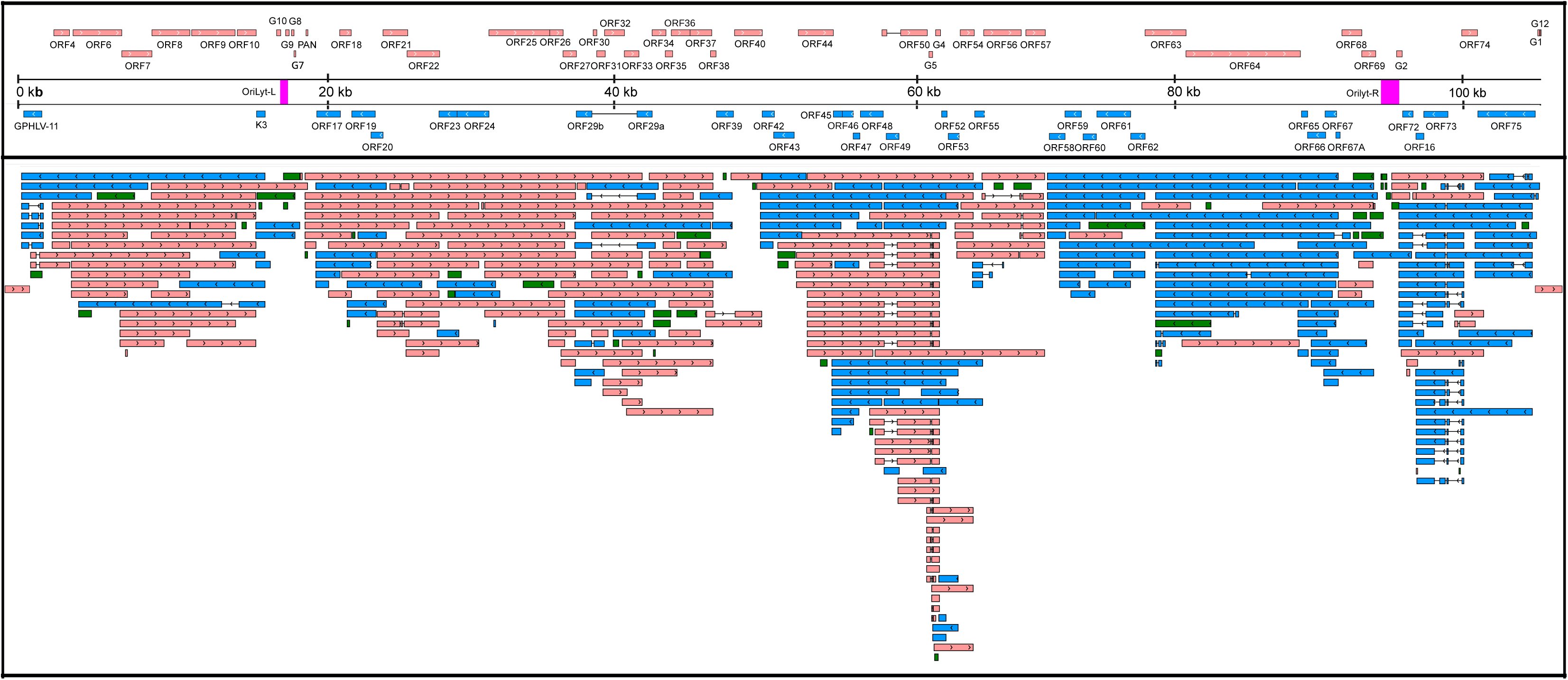
CaGHV-1 transcripts. This figure presents the canonical mRNAs and ncRNAs of CaGHV-1 along the reference genome. Transcripts with different splicing patterns are regarded as distinct canonical transcripts. Pink arrows indicate (+)-oriented RNAs, while blue arrows represent (-)-oriented RNAs. Additionally, antisense and replication-associated RNAs are shown in green.

We detected splicing events in both ncRNAs and mRNAs of CaGHV-1. In mRNAs, most introns were positioned in 5’-UTRs and 3’-UTRs. However, we identified introns in the coding regions of two genes (ORF50 and ORF57), where splicing resulted in different amino acid compositions at the N-terminal regions of the encoded proteins compared to the non-spliced transcripts. The intron structures of ORF50 and ORF57 are conserved and match those found in the homologous genes of KSHV. Furthermore, we found that ORF29 is composed of two exons, ORF29a and ORF29b, separated by a 3,093-nucleotide intron containing four genes (ORF30, ORF31, ORF32, and ORF33) oriented oppositely to the ORF29 gene. This intron arrangement is conserved in the homologous genes of PRV, HSV, KSHV, and EBV, except in alphaherpesviruses, where the intron contains only two genes (31–33). We identified several intron-containing ncRNAs in the G4-G5, ORF63-64, and OriLyt-R regions. The G4-G5 region, previously described only in KSHV, produces RNAs categorized as non-coding, despite containing small ORFs with no annotated function. In the ORF63-64 region, the antisense ncRNAs and the antisense segments of complex transcripts undergo splicing. Additionally, we detected an intron in a non-coding raRNA mapped to the OriLyt-R region **(Figure 4)**.

### Monocistronic viral mRNAs with canonical ORFs

In our analysis, we identified 54 canonical viral mRNAs, which were defined as the most abundant transcript variant for each viral gene (**Figure 4** and **Supplemental Table 5B).** This approach ensures that we capture the predominant functional outputs of the genome. Notably, one of the most prevalent viral transcript named ORF17.5 was located within ORF17 and featured a 5’-truncated ORF, exhibiting greater abundance than its full-length counterpart. The orthologous genes likewise express this embedded gene in all three herpesvirus subfamilies. Furthermore, we detected a transcript overlapping the genomic junction, beginning at the end of the G12 gene and ending within the GPHLV-11 gene.

### Multigenic CaGHV-1 transcripts

We identified 60 polycistronic viral mRNAs that encode two or more co-oriented ORFs and 108 complex transcripts containing two or more ORFs, with at least one positioned in an antiparallel orientation. The substantial number of complex viral transcripts indicates a high level of transcriptional complexity within the CaGHV-1 genome, primarily due to the presence of overlapping transcripts and alternative splicing events **(Figure 4** and **Supplemental Table 5B).** The longest polycistronic transcript measured 13,834 nucleotides, spanning seven genes within the ORF4-ORF10 region. While in KSHV, downstream genes within the ORF72-ORF71 and ORF35-36-37 transcripts have been shown to be translated via mechanisms such as termination-reinitiation, involving the utilization of upstream open reading frames (uORFs) (34), no similar mechanisms have been observed in other multigenic herpesviral transcripts. We detected three uORFs upstream of the ORF35 gene in CaGHV-1, but they are not as close to the ATG as in KSHV (**Supplemental Figure 1**). The longest complex transcript, measuring 22,936 nucleotides, shared a promoter with the PAN ncRNA and encompassed 16 genes spanning from ORF17 to ORF33.

### Non-coding viral transcripts

Non-coding RNAs comprise intergenic transcripts (limited to PAN), antisense transcripts, and likely numerous long 5’-UTR variants of mRNAs. A distinct class of ncRNAs is the replication origin-associated RNAs (raRNAs). Similar to KSHV, PAN is the most abundant ncRNA in the CaGHV-1 genome. Additionally, we identified 44 antisense RNAs (asRNAs), 33 of which are located within single genes, while 10 overlap two genes and 1 overlaps three genes (**Figure 4** and **Supplemental Table 5B**). For dRNA-Seq annotation, we utilized the NAGATA software developed by the Depledge laboratory (30). TATA boxes were identified in 26 antisense transcripts, 15 of which contained the TATTWAA sequence characteristic of late gene promoters in the KSHV virus. The average length of ncRNAs is 758.78 nt, with the shortest being 104 nt within ORF16 and the longest, 3,788 nt, located within ORF63-64. It is noteworthy that several asRNAs were detected in the ORF63-ORF64 and ORF75 regions, both in spliced and unspliced forms. Similar asRNAs have also been described in a closely related virus, Murine gammaherpesvirus 68 (35). Polycistronic transcripts with significant distances between their TESs and ATGs, as well as complex transcripts whose most upstream gene stands in an antiparallel orientation, are likely non-coding. We calculated the proportion of overlapping asRNA/mRNA pairs (**Supplemental Table 5B**).

Our analysis revealed numerous transcripts encoded in the vicinity of viral Oris, termed replication origin-associated RNAs, most of which are non-coding (**Figure 5)**. The replication origins of the CaGHV-1genome were identified by aligning it with the KSHV reference genome and mapping the corresponding KSHV replication regions onto the CaGHV-1 genome. Similar RNAs have also been previously described in other gammaherpesviruses, such as EBV and KSHV (36). We found that several transcripts, coterminating with K3 transcripts, overlap Orilyt-L with their 5’-UTRs. These transcripts are likely ncRNAs, given the large distance between their transcription and translation start sites. We also observed a very long RNA molecule in this region that fully encompasses the replication origin and co-terminates with PAN ncRNA. At the OriLyt-R, ORF72 produces a TES isoform that overlaps the replication origin with its 3’-UTR. Whether this transcript is involved in translation or serves solely to interfere with the replication process remains unknown. Additionally, multiple RNAs were identified with their promoter regions directly associated with OriLyt-R. A particularly intriguing discovery is that the transcripts originating from OriLyt-R utilize the TATTWAA promoter. Since these consensus sequences are recognized by LTF1 (encoded by ORF24), which facilitates the recruitment of RNA polymerase II, thereby regulating global transcription, we propose that this transcription factor may interfere with DNA replication by binding to the replication origin during the late stages of infection. Long complex RNAs were also detected at this region overlapping the entire lytic origin. We detected asRNAs overlapping ORF69, which is located near OriLyt-R but does not overlap it. Similar to CTO-S described in alphaherpesviruses (20, 37), which is also a non-overlapping ncRNA, these transcripts can be considered raRNAs. While we did not identify a latent replication origin in CaGHV-1, this does not rule out that it does not exist.

**Figure 5.**
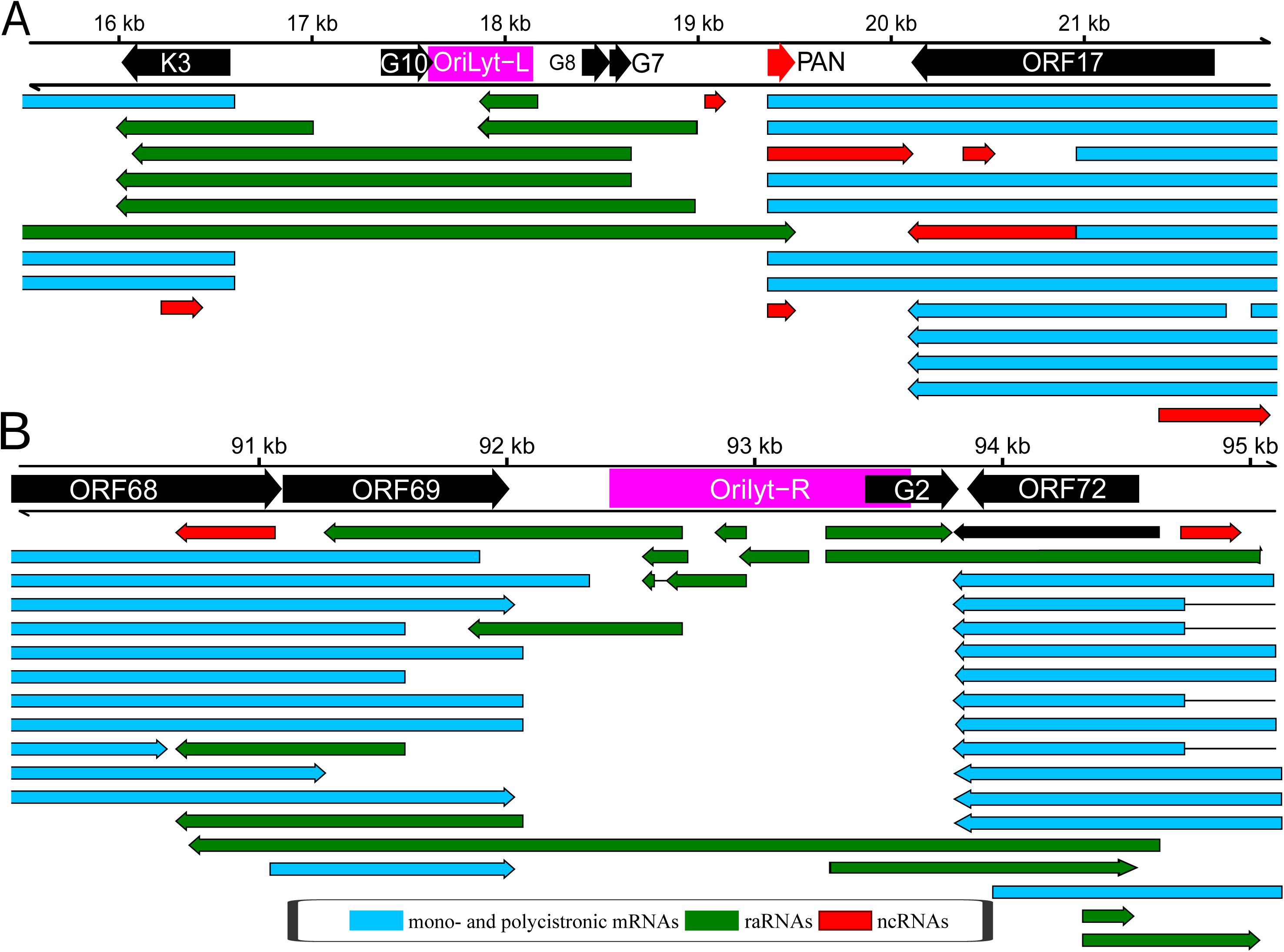
Replication origin-associated RNAs. **(A)** OriLyt-L: K3-PAN-ORF17 regions. **(B)** OriLyt-R: ORF69-ORF72 regions. Red arrows indicate non-coding RNAs, green arrows represent replication origin-associated RNAs (which can be either coding or non-coding), and blue arrows denote both monocistronic and polycistronic transcripts.

Additionally, we examined the potential of raRNAs to form RNA/RNA interactions using the IntaRNA program (38). Our analysis showed that three raRNAs (labeled 1, 2, and 3 in **Figure 4**) in the OriLyt-L regions slightly exceeded the required threshold (-30 kcal/mol) to be considered as potential interacting partners with the mRNAs of four viral genes: ORF9 (raRNA1, −3), ORF50 (raRNA1), ORF64 (raRNA1, −2, −3), and ORF73 (raRNA1, −3) **(Supplemental Table 5C)**. Further studies are needed to determine whether these high values indicate real functionality.

### Extensive genome-wide transcriptional overlaps among CaGHV-1 genes

In our work, we uncovered a remarkable level of transcriptional complexity, marked by transcriptional overlaps among convergent, divergent, and co-oriented genes in the CaGHV-1 genome (**Figure 6**). Our analysis showed that the entire viral genome is transcriptionally active on both DNA strands. Similar extensive transcriptional overlaps have been observed in other gammaherpesviruses, such as EBV (21 and 22) and KSHV (23). Notably, in the majority of convergent clusters (e.g., ORF18-ORF19, ORF22-ORF23, ORF27, ORF29b, ORF38-ORF39, ORF40-ORF42, ORF54-ORF55, ORF64-ORF65, G2-ORF72, ORF74-ORF75), we found ‘hard’ overlaps, where the 3’-ends of canonical transcripts overlap. In another group of convergent clusters (ORF10-K3, ORF44-ORF45, G4-ORF52, ORF57-ORF58), we observed ’soft’ overlaps, where canonical transcripts do not terminate within each other; however, transcriptional overlaps are occasionally generated through transcriptional readthrough between convergent or parallel-oriented gene pairs, or through head-to-head overlaps of divergent gene pairs.

**Figure 6.**
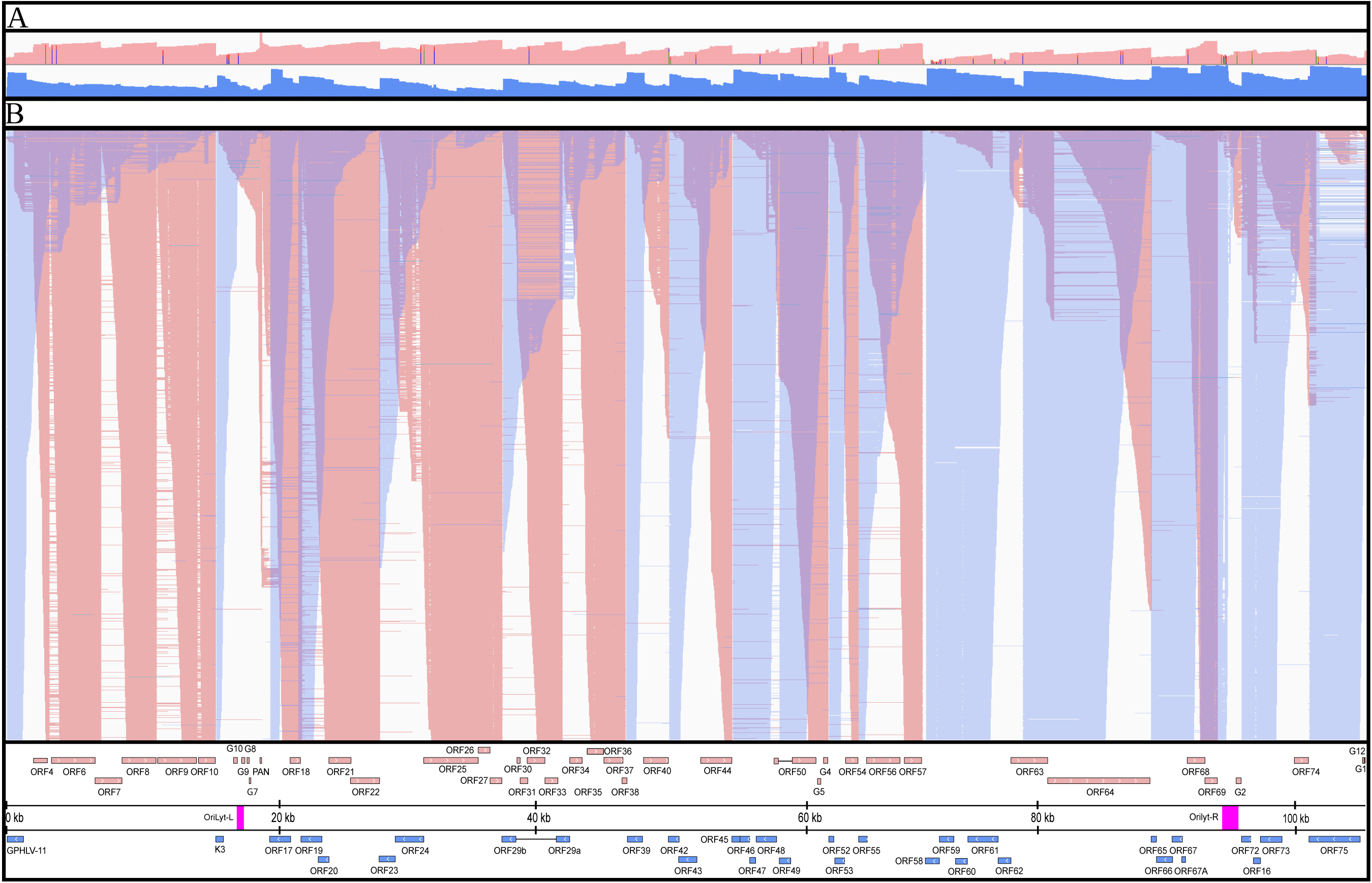
Overlaps of raw dRNA Reads. **(A)** The upper coverage plot shows the transcriptional activity of the viral genome, indicating that both DNA strands are transcriptionally active across the entire genome. Coverage values are plotted on a Log_10_ scale, with red representing the positive strand and blue representing the negative strand. **(B)** This panel highlights an extremely complex meshwork of transcriptional overlaps formed by genes arranged in head-to-head (divergent) and tail-to-tail (convergent) orientations. We hypothesize strong interference between the transcriptional machineries at the overlapping regions, which may represent a novel layer of gene regulation.

### Transcriptional Activity of ORF50 and Comparison with Gammaherpesviral Homologs

The homologs of the ORF50 gene product in gammaherpesviruses encode the replication and transcription activator protein (RTA), which is essential for driving the viral lytic cycle by activating the promoters of lytic genes, including that of ORF50. In our study, we found that the most abundant mRNAs of the CaGHV-1 ORF50 gene are initiated from two adjacent TSSs and contain four exons **(Figure 7A)**. The RTA protein consisting of 643 amino acid residues is encoded by the first two exons and is expressed as a 90-kDa protein **(Figure 7B)**. To assess the transcriptional activity of CaGHV-1 RTA (gpRTA) on a viral promoter, we cloned a 3-kb DNA region upstream of the ORF50 translational start site into a luciferase reporter vector. The luciferase reporter plasmid was co-transfected with increasing amounts of 3xFLAG-gpRTA into HEK293T and the guinea pig fibroblast cell line 104C1. Luciferase assays demonstrated that gpRTA greatly induced the ORF50 promoter in a dose-dependent manner in both cell lines **(Figure 7C)**. We also tested the effect of gpRTA on shorter 2 kb and 1 kb regions of the ORF50 promoter. We found that while gpRTA similarly activated the 3, 2, and 1 kb promoters of ORF50 in HEK293T cells, activation of the 2 kb promoter in the guinea pig cell line 104C1 was reduced by 4-fold compared to the 3 kb promoter, suggesting cell type and/or species-specific differences in the gpRTA-mediated promoter activation **(Figure 7D)**. To compare the promoter inducing function of gpRTA with its homologs from other gammaherpesviruses, we performed luciferase assays using the 3 kb promoter of CaGHV-1 ORF50 alongside RTAs from MHV68, EBV, HVS, KSHV, CaGHV-1, and RRV **(Figure 7E)**. The results revealed that while gpRTA efficiently induced the CaGHV-1 ORF50 promoter in both human and guinea pig cell lines, the other gammaherpesvirus RTAs showed varying levels of promoter activation in the two cell lines **(Figure 7F)**. In conclusion, our findings indicate that gpRTA shares functional similarities with its gammaherpesvirus homologs regarding its ability to induce the promoter of its own gene.

**Figure 7.**
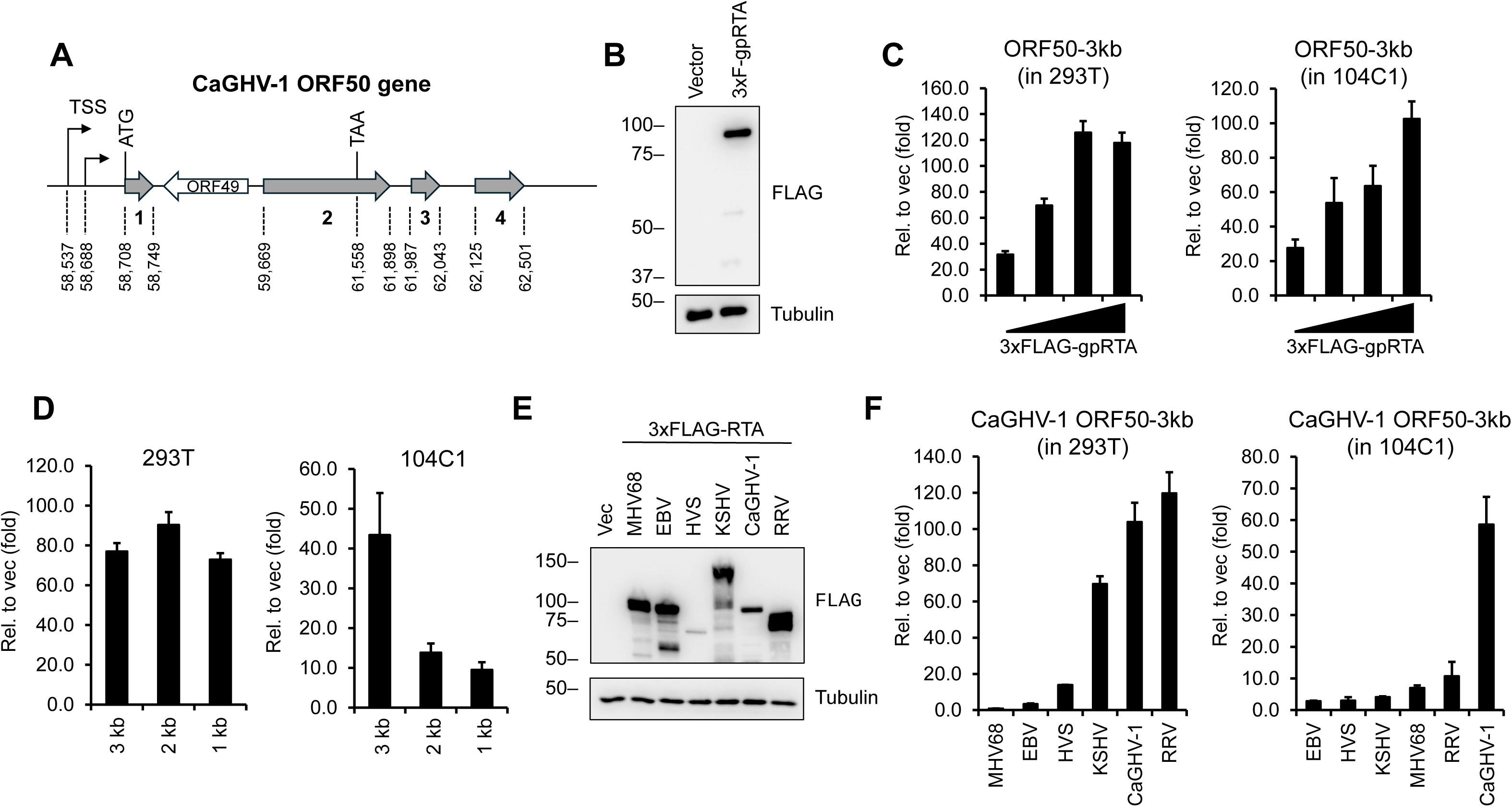
Evaluating the transcriptional activity of CaGHV-1 RTA. **(A)** Gene structure of ORF50 encoding gpRTA in the CaGHV-1 genome. The ORF50 gene has 4 exons. The genomic coordinates are based on OQ679822.1 (GenBank). **(B)** Protein expression of N-terminally 3xFLAG tagged CaGHV-1 RTA in transfected 293T cells. **(C)** Testing the inducibility of the 3 kb promoter region of CaGHV-1 ORF50 by CaGHV-1 RTA in 293T and 104C1 cell lines using luciferase reporter assays. **(D)** Measuring the inducibility of the ORF50 promoter with differing lengths by ORF50 promoter in luciferase reporter assays. **(E)** Western blot showing the protein expression of N-terminally 3xFLAG tagged RTAs derived from the indicated gammaherpesviruses. **(F)** Analyzing the transcriptional activity of RTAs derived from the indicated gammaherpesviruses on the 3 kb promoter region of CaGHV-1 ORF50 in 293T and 104C1 cells.

## DISCUSSION

In this study, we integrated ONT dcDNA-Seq and dRNA-Seq data and used two bioinformatics software tools (LoRTIA and NAGATA) to construct a comprehensive transcriptome map of CaGHV-1. Specifically, we mapped a number of canonical mono- and polycistronic mRNAs, along with non-coding transcripts, including intergenic (PAN), antisense, and replication origin-associated RNA molecules, as well as transcripts with truncated ORFs. Furthermore, we also described fusion and complex RNA molecules. We also identified cis-regulatory elements with single-nucleotide resolution, including TSS initiation motifs, promoter elements, poly(A) signals, 3’-cleavage sites, and splice junctions. For TSS identification, we employed the LoRTIA software package (24), which enabled us to filter out false 5’-ends by excluding reads with incorrect template switching adapters. To reduce false positives, only TSS present in all three samples were accepted, leading to the identification of 165 TSS and 93 TATA-boxes. Notably, the TATTWAA motif - previously described in KSHV, EBV, and CMV - was detected upstream of several TSSs, distinct from the TATA-box motif used by early genes (39). These sequences serve as binding sites for LTF1 and vTA, which are crucial for the transcription of late herpesviral genes, as they recruit RNA Polymerase II (29). Moreover, the InR pattern we observed aligns with previous findings across all herpesvirus families, indicating that the nucleotide composition around the TSS is highly conserved (13, 22, 26).

To identify mRNA 3’-ends, we employed the LoRTIA software package, selecting only those 3’- ends that were present in all three samples and contained three or more adenines at the 3’-end. This approach is currently the most accurate method for detecting mRNA 3’-ends, as it effectively filters out false 3’-ends caused by false priming or template switching (24). Additionally, we used dRNA-Seq for validation, which further eliminates false results because it processes native RNA molecules. Through this process, we identified 143 TESs, most of which contained the canonical AAUAAA motif characteristic of eukaryotic mRNAs. This motif has been detected across all three herpesvirus subfamilies, suggesting that efficient 3’-end cleavage is regulated by host factors (40, 41).

The presence of RNAs overlapping the lytic replication origin has been observed in several DNA viruses (36). In HCMV, the RNA 4.9 is the most abundant viral RNA and plays a regulatory role in replication, while in KSHV, expression of the T1.5 transcript is associated with various biological functions, such as promoting cell survival, immune modulation, and contributing to angiogenesis and pathogenesis in infected cells (42–44). In this study, we identified several raRNAs in both lytic Ori regions of CaGHV-1. Some of these RNAs originate directly from the Ori, while others, including relative short ncRNAs and long polycistronic transcripts, overlap with the Ori and extend into the cis-regulatory elements of neighboring genes. One notable overlap spans from ORF8 to PAN. It is intriguing to speculate why this long transcript uses the PAN PAS signal rather than terminating earlier. One possible explanation is that during the transcription of long RNAs, collisions between RNA and DNA polymerases may occur. Additionally, such overlaps could inhibit the transcription of genes, thereby helping to separate the processes of replication and transcription (45). Thus, these RNAs may play multiple roles in regulating the viral life cycle. Moreover, an interesting observation is that the OriLyt-R region contains the TATTWAA motif necessary for late gene transcription, hinting at an interaction between transcription regulation and DNA replication. Understanding this phenomenon could pave the way for new research directions, potentially leading to interventions targeting viral molecular mechanisms.

Using dRNA-Seq, validated by dcDNA-Seq and parallel sequencing of mpox transcripts (which lack introns), we identified numerous introns in both the coding and UTR regions of several mRNAs, as well as in non-coding transcripts of CaGHV-1. For spliced transcripts, we observed a pattern similar to that found in KSHV and MHV68. Notable examples include the ORF50 (RTA) mRNA, which plays a key role in initiating and regulating the lytic cycle of the virus, and the ORF64 mRNA (35, 46, 47). Additionally, we found a high degree of isoform diversity in ORF73 (LANA), a protein involved in maintaining latent infection and immune evasion (48).

We also detected a large number of antisense, complex, and polycistronic transcripts, contributing to an intricate network of gene overlaps throughout the genome. These overlaps occur between convergent, divergent, and parallel genes. Read-throughs between convergent genes and overlaps between divergent genes result in antisense segments on the generated transcript. In HSV-1, it has been demonstrated that these form dsRNAs, which are inhibited by the virion host-shutoff (VHS) gene product, suggesting that the actual frequency of read-throughs may be higher than observed (49). Genome-wide transcriptional read-throughs contribute to widespread antisense activity, consistent with findings in other gammaherpesviruses. We have proposed a "transcriptional interference network" hypothesis for this phenomenon, suggesting that competition and collision between transcriptional machineries during interactions between neighboring genes serve as a regulatory mechanism (50). Moreover, transcriptional overlaps were detected not only between genes but also at the genomic ends, where RNAs spanning the circular genomic junctions were observed. These RNA molecules have also been confirmed in KSHV and EBV (51); however, they are often overlooked in studies due to their low abundance (52).

Based on the similarity in gene organization between KSHV and CaGHV-1, we cloned the CaGHV-1 ORF50 gene, which is anticipated to encode RTA, a viral transcription factor conserved across all gammaherpesviruses and essential for inducing the lytic cycle (53). A key feature of gammaherpesviral RTAs is their ability to bind to and activate the ORF50 promoter, initiating a positive feedback loop that enhances the expression of RTA and other lytic genes. Our findings show that CaGHV-1 RTA strongly activates the CaGHV-1 ORF50 promoter. Interestingly, several homologs of CaGHV-1 RTA from other gammaherpesviruses also activate the CaGHV-1 ORF50 promoter, highlighting functional similarities between CaGHV-1 RTA and other gammaherpesvirus RTA proteins in promoting viral transcription. Further studies are needed to assess the extent to which CaGHV-1 RTA shares functional similarities with human gammaherpesvirus RTAs, particularly in regulating viral and host gene expression and influencing protein degradation. By mapping the genes and regulatory regions of CaGHV-1, and cloning its RTA, we pave the way for developing CaGHV-1 as a novel model to investigate the biology and pathogenesis of human gammaherpesvirus infections.

## MATERIALS AND METHODS

### Cells and Virus

CaGHV-1 (VR-543), the guinea pig (GP) fibroblast cell line 104C1 (CRL-1405), and HEK293T cells were purchased from ATCC. The cell lines 104C1 and HEK293T were grown in RPMI-1640 and DMEM media, respectively, supplemented with 10% FBS and penicillin/streptomycin. CaGHV-1 was amplified in the cell line 104C1 followed by the concentration of the virus supernatant by ultracentrifugation. 10⁴ GP cells were infected with CaGHV-1, and the cells were collected at eight time points (4h, 8h, 16h, 24h, 48h, 72h, 96h, and 120h) post-infection. The samples from each time point were mixed in equal volumes for both dRNA-Seq and cDNA-Seq.

### DNA cloning and luciferase reporter assay

The protein coding sequence of CaGHV-1 ORF50 was PCR amplified and cloned into the pCDH-CMV-MCS-EF1-puro expression vector using In-Fusion cloning (TaKaRa). The cloning of the other gammaherpesvirus RTAs has been published in our previous study (54). CaGHV-1 RTA was expressed as an N-terminally 3xFLAG-tagged protein in HEK293T cells by transfecting the cells with PEI (Polysciences). The CaGHV-1 ORF50 promoter fragments were PCR amplified and cloned into pGL4.15 luciferase reporter vector (Promega) using In-Fusion cloning. In the luciferase reporter assays, 100 ng of reporter plasmids were co-transfected with 400 ng of RTA expression plasmids. To assess the effect of varying RTA levels, increasing amounts of RTA expression plasmids (50, 100, 200, and 400 ng) were used for transfection. The luciferase assay was performed as described previously (55).

### Isolation of RNA

Total RNA was isolated using TRIzol reagent (Invitrogen) according to the manufacturer’s protocol with some modifications. After adding chloroform to the cells lysed in TRIzol, the lysates were spun down. Afterwards the supernatants were mixed with 100% ethanol and added to RNeasy columns. The RNA purification was performed by the protocol of the RNeasy kit (Qiagen). The polyadenylated RNA enrichment was carried out using Lexogen’s Poly(A) RNA Selection Kit V1.5. The RNA samples were bound to beads, washed, and hybridized. After incubation and washing, the polyadenylated RNA was eluted in nuclease-free water and stored at -80°C for subsequent analysis.

### Direct cDNA sequencing

Direct cDNA sequencing was performed on the Oxford Nanopore Technologies (ONT) Mk1B and Promethion P2 Solo devices. For the preparation of sequencing libraries, we used the Ligation Sequencing V14 – dcDNA-Seq (SQK-LSK114) kit. Currently, the manufacturer does not provide barcoding for direct cDNA libraries, so we combined it with the Ligation Sequencing gDNA - Native Barcoding Kit 24 V14 (SQK-NBD114.24) for sample barcoding. For each sample, the initial amount was 1 µg total RNA, which was mixed with a VN primer (User-Supplied VNP 2 µM, ordered from IDT) and a 10 mM dNTP mix, then incubated at 65°C for 5 minutes. This was followed by cooling on a pre-chilled freezer block for 1 minute, and then adding the 5x RT Buffer, RNaseOUT (Thermo Fisher Scientific), and the Strand-Switching Primer (User-Supplied SSP 10 µM, ordered from IDT), followed by heating at 42°C for 2 minutes.

Reverse transcription and first-strand cDNA synthesis were carried out using the Maxima H Minus Reverse Transcriptase enzyme (Thermo Fisher Scientific), with the reaction occurring at 42°C for 90 minutes and enzyme inactivation at 85°C for 5 minutes. The RNA molecules were digested from the RNA-cDNA hybrids using RNase Cocktail Enzyme Mix (Thermo Fisher Scientific) at 37°C for 10 minutes. For second-strand cDNA synthesis, we used LongAmp Taq Master Mix [New England Biolabs (NEB)] and a PR2 Primer (User-Supplied 10 µM, ordered from IDT), with the PCR reaction involving Denaturation at 94°C for 1 minute (1 cycle), Annealing at 50°C for 1 minute (1 cycle), and Extension at 65°C for 15 minutes (1 cycle).

The double-stranded cDNAs then underwent end-repair and dA-tailing using the NEBNext® Ultra II End Repair/dA-tailing Module, incubated at 20°C for 5 minutes and 65°C for 5 minutes. For subsequent steps, we used the ONT Ligation Sequencing gDNA - Native Barcoding Kit 24 V14 (SQK-NBD114.24) protocol for sample barcoding. End-prepped DNAs were barcoded, and NEB Blunt/TA Ligase Master Mix (NEB) was added, followed by a 20-minute incubation at room temperature (RT), then EDTA addition. This was followed by ligation of the Native Adapter (NA) included in the kit, using the NEBNext Quick Ligation Module (NEB) enzyme and buffer. AMPure XP Beads (AXP, from the ONT kit) were used for DNA purification after each enzymatic step. Samples were then eluted in nuclease-free water. For concentration measurement, we used the Qubit 4.0 fluorometer and the Qubit dsDNA HS Assay kit. From the prepared cDNA libraries, 50 fmol/flow cell was loaded into R10.4.1 flow cells. To prevent "barcode hopping," early time points were sequenced separately using an R10.4.1 flow cell (FLO-MIN114) and an R10.4.1 flow cell (FLO-PRO114M), and later time points were sequenced using an R10.4.1 flow cell (FLO- PRO114M).

### Native RNA sequencing

For native RNA sequencing, we used the ONT Direct RNA Sequencing (SQK-RNA004) kit. For library preparation, we pooled 1 µg of total RNA from the samples into 8.5 µl. First, we ligated an RT Adapter (RTA) to the samples using NEBNext® Quick Ligation Reaction Buffer (NEB), T4 DNA Ligase (2M U/ml, NEB), and RNaseOUT™ Recombinant Ribonuclease Inhibitor (Invitrogen), followed by a 10-minute incubation at room temperature. The next step involved adding a reverse transcription master mix to the adapter-ligated RNA, which contained 10 mM dNTPs, 5X First-Strand Buffer, and DTT. Synthesis of the cDNA strand was performed using SuperScript™ III Reverse Transcriptase (Thermo Fisher Scientific), with the reaction run at 50°C for 50 minutes, followed by inactivation at 70°C for 10 minutes. The RNA-cDNA hybrids then had the RNA Ligation Adapter (RLA) ligated using NEBNext Quick Ligation Reaction Buffer and T4 DNA Ligase. After each enzymatic reaction, we used Agencourt RNAClean XP beads for purification. For concentration measurement, we used the Qubit 4.0 fluorometer and Qubit dsDNA HS Assay Kit. The prepared library was sequenced on the Promethion P2 Solo device using an RNA flow cell (FLO-PRO004RA).

### Bioinformatics

The raw current signals underlying the analyses were initially assigned to nucleotides using the Dorado-0.8.2 basecaller. Reads were aligned to the reference genome (accession number: OQ679822.1) using the minimap2 software with the following parameters: Y -C5 -ax splice –cs. The identifiers and their availability in the European Nucleotide Archive (ENA) database are listed in **Supplemental Table 6**. SeqTools (https://github.com/moldovannorbert/seqtools) was employed for promoter element identification and basic statistical calculations. The LoRTIA tool, developed by our research group, was used to detect TSS, TES, and introns ("features") and to annotate transcripts (https://github.com/zsolt-balazs/LoRTIA, v0.9.9). The first phase of the process involved identifying sequencing adapters, homopolymer As, and removing erroneous reads generated by RNA degradation, template switching, or faulty priming. The parameters for this step were as follows: Samprocessor.py --five_adapter GCTGATATTGCTGGG --five_score 14 --check_in_soft 15 --three_adapter AAAAAAAAAAAAAAA --three_score 14 input output.

In the next step, potential TSS and TES positions were identified. The first nucleotide that did not align with the adapter was marked as a potential TSS, while the last nucleotide that did not align with the homopolymer A was designated as a potential TES. This analysis was conducted for each ’sam’ file using the following commands: Stats.py -r genome -f r5 -b 10 and Stats.py -r genome -f l5 -b 10 for TSS detection, Stats.py -r genome -f r3 -b 10 for TES detection, and Stats.py -r genome -f in for intron identification. Adapter alignment was evaluated using the Smith-Waterman algorithm. False 3’-ends arising from false priming or template switching were removed if at least three adenines preceded the homopolymer A. To further validate the TES positions and exclude those resulting from internal priming or other errors, the poly(A) length estimation module in the Dorado software package was used to identify and estimate poly(A) sites in dRNA-Seq samples (default settings were applied). Reads in the dRNA samples were identified and assigned to transcripts using the NAGATA software (default settings) (30).

In the third phase, potential TSSs and TESs were evaluated using the Poisson distribution to filter out random start and end positions caused by RNA degradation. Significance was corrected using the Bonferroni method. Features observed in fewer than two reads or with coverage of less than 1% were excluded from further analysis. Additional criteria required TSSs to appear in at least three direct cDNA samples, and TESs to occur in direct RNA samples. The command Gff_creator.py -s poisson -o was used for this step. Subsequently, we ran the transcript annotation module, which assigns validated features (TSSs, TESs, and introns) to each read using the parameters: Transcript_Annotator_two_wobbles.py -z 20 -a 10.

Statistical charts in **Figure 1**, along with nucleotide distribution and Log₁₀ line plot diagrams in **Figures 2 and 3**, were visualized using the Matplotlib Python library. Nucleotide sequences were extracted using the Bedtools getfasta software package. The Integrative Genomics Viewer (IGV) was used for overall transcriptome visualization.

## Data availability

The sequencing datasets generated in this study are available at the European Nucleotide Archive under the accession: PRJEB80811 and link https://www.ebi.ac.uk/ena/browser/view/PRJEB80811.

## Acknowledgements

The research was funded by the National Research, Development and Innovation Office (NRDIO), through the Researcher-initiated research projects (Grant number: K 142674) awarded to ZB. ZT was supported by the NIH grant R01DE028331. DT was supported by NRDIO FK 142676. The publication fee was covered by the University of Szeged, Open Access Fund: 7358.

## Ethics declarations

Not applicable

### Conflicts of interests

The authors do not declare any conflicts of interest.

### Author contributions

**G.T:** carried out bioinformatic analyses, visualization, and drafted the manuscript

**Á.D:** participated in long-read sequencing

**Á.F:** contributed to bioinformatics and visualization

**D.T:** contributed to library preparation, participated in data interpretation, and drafted the manuscript

**M.M**: contributed to library preparation

**S.L:** cultivated the cells and prepared RNA samples

**A.M.P:** did the DNA cloning and the promoter reporter assays

**Z.T:** contributed to the experiment design and drafted the manuscript

**Z.B:** conceived and designed the experiments, supervised the study, and wrote the manuscript

All authors read and approved the final paper.

## Supplemental Figure and Tables

**Supplemental Figure 1.** Usptream ORFs in the ORF35 transcript

**Supplemental Table 1.** Statistics of the Oxford Nanopore PromethION dcDNA and dRNA sequencing.

**Supplemental Table 2.** Positions of the TSSs, their sample-specific abundances, and associated promoter elements.

**Supplemental Table 3.** Positions of the TESs, their abundances in each sample, and the associated poly(A) signals.

**Supplemental Table 4.** The positions of introns and their abundances across different samples.

**Supplemental Table 5.** List of spliced (A), non-spliced (B) transcripts detected by LoRTIA, and replication-associated RNAs (C) including their binding energies and target genes.

**Supplemental Table 6.** Access identifiers for fastq files generated by long-read sequencing, available in the European Nucleotide Archive (ENA) database.

